# Bacterial communities in residential wastewater treatment plants are physiologically adapted to high concentrations of quaternary ammonium compounds

**DOI:** 10.1101/2023.08.01.551460

**Authors:** Luz Chacon, Keilor Rojas-Jimenez, Maria Arias-Andres

**Affiliations:** Instituto de Investigaciones en Salud (INISA), Universidad de Costa Rica, San José, Costa Rica; Escuela de Biología, Universidad de Costa Rica, 11501-2060 San José, Costa Rica; Instituto Regional de Estudios en Sustancias Tóxicas (IRET), Universidad Nacional, Campus Omar Dengo, Apdo 86-3000, Heredia, Costa Rica

**Keywords:** Activated sludge, Benzalkonium chloride, EcoPlate System, Functional traits, *Pseudomonas*

## Abstract

Benzalkonium chloride (BAC) is a quaternary ammonium compound (QAC) widely used as the active ingredient of disinfectants. Its excessive discharge in wastewater is constant and in high concentrations, likely affecting the physiology of microbial communities. We compared the community physiological profile of activated sludge bacteria with and without *in vitro* previous exposure to a high concentration of BAC (10 mg/L). We measured the community functional diversity (FD), carbon substrate multifunctionality (MF), and the median effective concentration that inhibits carbon respiration (EC_50_) using Biolog® EcoPlatesTM supplemented with a gradient of 0 to 50 mg/L of BAC. Surprisingly, we did not find significant differences in the physiological parameters between treatments. Certain abundant bacteria, including Pseudomonas, could explain the community’s tolerance to high concentrations of BAC. We suggest that bacterial communities in wastewater treatment facilities’ activated sludge (AS) are “naturally” adapted to BAC due to frequent and high-dose exposure. We highlight the need to understand better the effects of QACs in wastewater, their impact on the selection of tolerant groups, and the alteration in community metabolic profiles.

## 1. Introduction

Quaternary ammonium compounds (QAC) are a diverse group of substances that find widespread use in a variety of applications including domestic, agricultural, healthcare, and industrial settings. QACs function as surfactants, emulsifiers, fabric softeners, pesticides, corrosion inhibitors, paint additives, cosmetics, personal care products, and disinfectants [1]. Benzalkonium chloride (BAC) is one of the most commonly QACs used in the disinfectant industry. BAC is typically sold as a mixture of different alkyl chain lengths ranging from C8 to C18, C12 and C14 derivates exhibiting higher biocidal activity [2]. BAC action mode involves the ability of the alkyl chain to permeate cell membranes and disrupt their physical and biochemical properties, while the remaining portion of the molecule remains at the membrane, disrupting the ion charge distribution [3].

BAC is effective against a wide range of microorganisms including yeast, fungi, gram-positive and gram-negative bacteria, and certain viruses [4]. With the onset of SARS-CoV-2 pandemic, the use and disposal of QACs in urban wastewater have increased dramatically. In the United States, over one million pounds (543 000 kg) of BAC were manufactured or imported prior to 2020. As a result, the global disinfectant market is projected to grow by over 6% from 2016, reaching over $8 billion by 2021 [5]. In addition, some governments have approved using new cleaning products for virus destruction. Up to 50% of approved products contain QAC; with 95% of them utilizing BAC as the primary active component [6].

BAC is biodegradable under aerobic conditions, and its environmental concentrations fluctuate, exposing microbial communities from non-inhibitory to over-inhibitory concentrations [7]. The bacterial consortium involved in BAC biodegradation in sewage and activated sludge (AS) primarly comprises Proteobacteria such as *Pseudomonas, Achromobacter*, and probably other uncultured microorganisms [8]. However, some studies have shown that BAC use can lead to the development of antibiotic resistance in bacterial communities. [9]. Moreover, BAC concentrations of 2 mg/L or higher in wastewater treatment systems can inhibit the nitrification and biological nutrient removal processes [10].

Molecular ecology tools are commoly used to monitor the adaptation of microbial communities and changes in antibiotic-related resistance genes. However, it is also important to use functional methods that measure activity to assess the effects of contaminants and the tolerance of communities. One practical, low-cost, and effective screening method to assess the effects on the functional diversity of AS microbes in wastewater treatment facilities (WWTPs) at community level is the Biolog® EcoPlate™ system [11]. EcoPlate™ is a helpful approach to evaluate the carbon sources used by a microbial community, which uses a redox reaction to detect carbon source metabolization by each microbial community. The rate or the color development variance depends on the specific microbial community, and this pattern is referred to as the “community-level physiological profile” [12].

In a previous study we demonstrated that AS bacteria from a WWTP in Costa Rica exhibited high resilience to changes in community structure while increasing the abundance of genes associated with QACs resistance when exposed to a pulse of BAC in the range of 10 mg/L [13]. In this article, we investigate the effects of a second exposure to increasing BAC concentrations on microbial activity in both BAC-exposed and non-exposed communities using Biolog® EcoPlates™. We also analyze the taxonomic composition of the bacterial community present in the EcoPlates™ to gain insight into wich part of the community is represented in the microbial activity measurements. Our hypothesis is that the frequent domestic use and disposal of disinfectants containing BAC leads to the adaptation of bacterial communities in AS to concentrations in the range of mg/L of BAC, making *in vitro* exposure likely to cause only slight changes in the bacterial composition in the EcoPlates™. The increasing use of disinfectants containing compounds like BAC does not go unnoticed in the environment. Therefore, this study aims to shed light on the possible permanent impacts of BAC-related compounds on the physiology and taxonomic composition of bacterial communities in WWTPs.

## 2. Materials and Methods

### 2.1. Sample preparation

A small residential activated sludge plant, representing the most common type WWTP in Costa Rica, was selected. The sewage system serves a population of 976 and is located in the Costa Rican Central Valley (9°55’14” N, 84°14’34’’W) at an elevation of 1,400 meters above sea level with an average temperature of 22 °C. Four liters of AS were collected from the aeration tank (4 meters deep) using a metal bucket approximately 50 cm deep. The pH, dissolved oxygen, conductivity, and temperature of the tank were recorded in the AS sample with a HACH® HQ40d portable meter. The sample was transferred to a sterile amber glass bottle container, kept at 4 °C, and immediately transported to the laboratory. Once in the laboratory, the sample was divided into two portions of 2 L each and stored for 12 hours at 25 ± 2 °C with continuous airflow. For the enrichment test, one portion of the sample was dosed with 10 mg/L of benzalkonium chloride (BAC, ≥95.0% Fluka 12060, Sigma Aldrich), divided into three replicas of 0.15 L, and incubated at 25 ± 2 °C, with agitation (150 rpm) and aeration for 96 hours. A non-BAC enriched control sample was treated with the same conditions. A summary of the workflow is presented in Figure 1.

**Figure 1.**
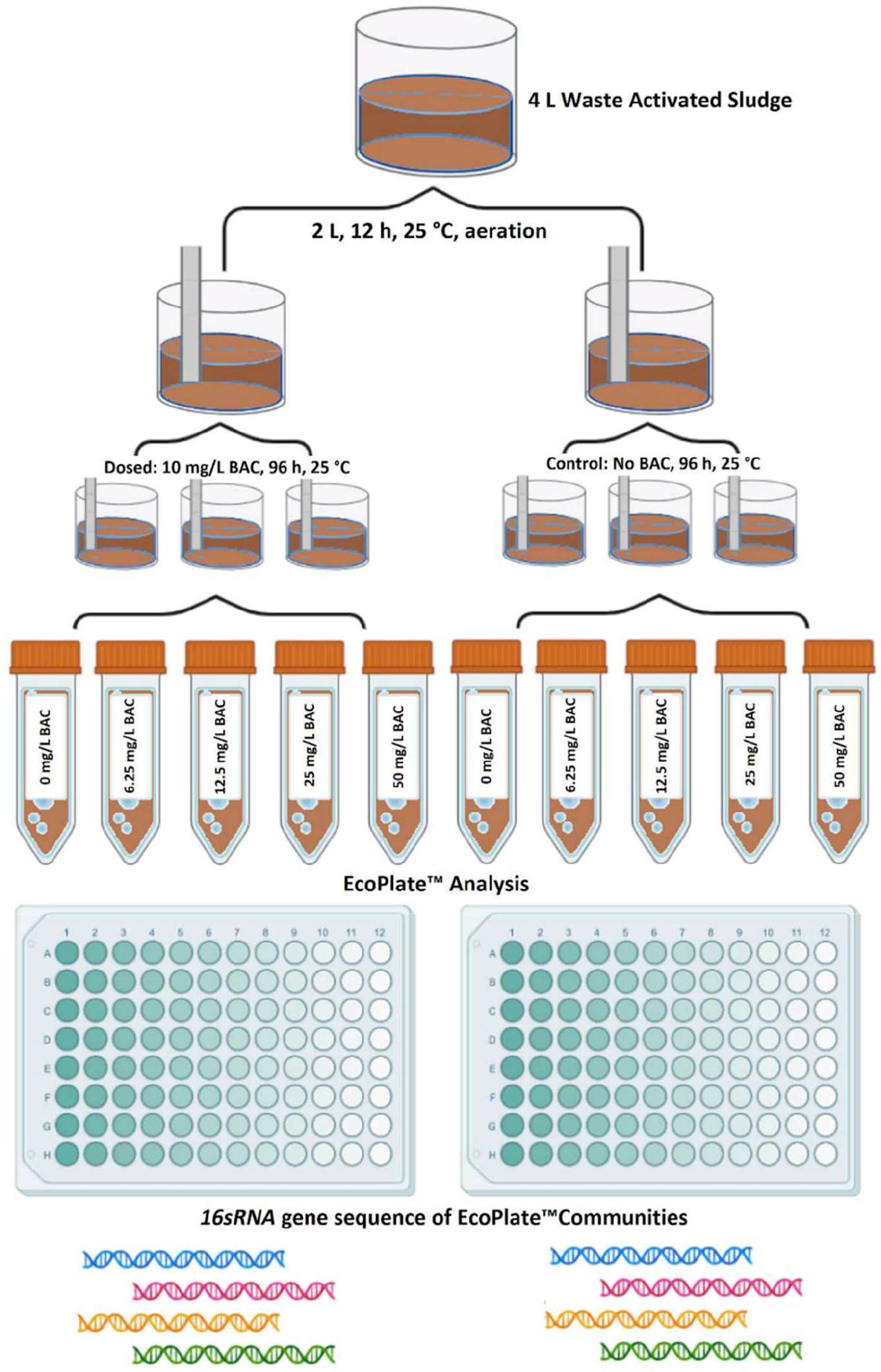
General methodology workflow. Figure created with BioRender.

### 2.2. Benzalkonium chloride quantification

BAC concentration was determined at three time points: 1. Before lab storage, 2. Immediately after enrichment pretreatment, and 3. Before EcoPlate™ inoculation. The method for BAC extraction from AS, followed Chacón et al, 2021 [14] using organic solvents and a rotary evaporator system. BAC determination was performed using an Acquity Ultra Performance Liquid Chromatography system (UPLC), consisting of a Waters Acquity binary solvent manager, autosampler, and Photodiode Array Detector (PDA) coupled with a Quadrupole Time of Flight (Q-ToF) (Waters Synapt G1), (Waters Corp., Milford, MA, USA) in series. The samples were analyzed in triplicate and the three main BAC homologs were identified based on their retention time, and high-resolution molecular mass determination (M-H)+ comparisons with reference standards.

### 2.3. Physiological Profile determination with EcoPlates™

We used EcoPlates™ (Biolog®, USA) to analyze the carbon substrate utilization by a microbial community after a pulse exposure to BAC. Briefly, EcoPlates™ are 96-well plates containing three repeated sets of 31 carbon sources and employ a tetrazolium redox dye as an indicator of microbial metabolism. The tetrazolium is reduced to visible purple when microorganisms utilize each carbon source at a constant temperature, and redox reaction can be quantified using spectrophotometric techniques. For the experiment, we prepared seven individual 10^−1^ dilutions of AS (enriched and control) using 10 mM phosphate buffer (pH=7.0); each dilution was mixed with a gradient of BAC (0 mg/L, 6.25 mg/L, 12.5 mg/L, 25 mg/L, 50 mg/L). We inoculated 100 μL of each preparation into each well and incubated it in the dark at 25 ± 2 °C with >85% humidity. Absorbance at 595 nm was diary measured for seven days [15]. For each substrate, the concentration of the BAC that caused an inhibitory effect (EC_50_) was estimated using the area under the curve (AUC) of the replicates of each BAC concentration for both the BAC enriched and control groups. A log-logistic regression based on all measurements was used to estimate EC_50_.

### 2.4. DNA extraction and sequencing

Upon completion of the physiological profile determination, the microorganisms grown on the EcoPlates™ were recovered and divided into nine pools corresponding to control (0 mg/L of BAC), and concentrations of 6.25 mg/L, 12.5 mg/L, 25 mg/L, 50 mg/L for both enriched and non-BAC’s enriched pretreatments. Each pool was transferred to a15 ml plastic tube, centrifugated, and the pellets were sujected to DNA extraction. The DNA extraction procedure involved adding extraction buffer (100 mM Tris, 100 mM EDTA, 200mM NaCl, 1% PVP, 2% CTAB), proteinase K (0.3 mg, Invitrogen®, USA), and lysozyme (0.5 mg, Invitrogen®, USA), followed by a 2-hour incubation. Chloroform-alcohol (v/V 24:1) was added, two wash steps were made with isopropanol, and a final washing step was conducted with ethanol 70%; after that, the clean DNA was resuspended in ultrapure water. The quality of the extracted DNA was assessed spectrophotometrically (Nanodrop2000, Thermo, USA). TheV3-V4 region of *16SrRNA* gene was amplified using 341F (5’ CCTACGGGNGGCWGCAG 3’) and 785R (5’ GACTACHVGGGTATCTAATCC 3’) primers [16], and the samples were sequenced using MySeq PE300 (Illumina®, USA) by MR DNA (www.mrdnalab.com, Shallowater, TX, USA).

### 2.5. Statistical and data analysis

#### 2.5.1. Physiological Data Preparation

First, to obtain the area under the curve (AUC) for each subtrate, the OD595 nm values were integrated. The average AUC value was calculated for each EcoPlate™. Next, the AUC values were normalized by subtracting the AUC from the “water” control (no substrate) and dividing it into the total EcoPlate™ reading period (7 days) [17]. For each BAC treatment (Control or BAC) and concentration on the EcoPlate™ (0,6.25, 12.5, 25, 50 mg/L), we computed the richness of functions and determined the functional diversity profile, following the statistical approach described by Miki et al. [18]. Additionally, we used the dose-response approach of EcoPlate™ inoculation and BAC supplementation to calculate the BAC concentration inhibiting 50% of substrate utilization in each Treatment (EC_50_).

#### 2.5.2. Multifunctionality (MF) Index and Diversity Profile (FD) Calculation

We calculated the multifunctionality index to evaluate the richness of functions (MF=number of substrates utilized). To classify the AUC measured in the EcoPlate™ as the presence (1) or absence (0) of carbon substrate metabolism, we used a quantile-thresholding method [19]. A matrix of MF values was obtained for each sample (Treatment+BAC concentration on EcoPlate™) and threshold (0.1 to 0.9). All the statistical analyses were performed in R [20]. Initially, the MF indexes of Control and BAC per threshold were compared using the Kruskal-Wallis test (KW). Then, samples (Treatment+BAC concentration on EcoPlate™) were compared by KW, and multiple comparisons with Dunn’s Test at alpha = 0.05 were conducted for thresholds 0.1, 0.5, and 0.9). The MF values obtained per sample from all thresholds were also compared using the KW test. A dissimilarity matrix for each sample was created from the AUC measurements of the EcoPlate™, using a Bray-Curtis method to analyze the functional diversity profiles (FD) [21]. The matrices were compared using permutational multivariate analysis of variance (Permanova) at a probability value of 0.05 and with 999 permutations, using the function adonis from the “vegan” package [22]. The p-values from pairwise comparisons were adjusted according to the Benjamini-Hochberg method. The dissimilarity matrices were visualized in a non-metric multidimensional scaling plot using “ggplot2” package [23]. EC_50_ values were determined with package “drc” using the function “mselect()”to select the appropriate model for each dose∼response relation (per substrate). An analysis of variance and model fit were then performed using functions “anova()” and “modelFit()” respectively [24]. Finally, EC_50_ values were compared between control, BAC-treated AS, and both; and per individual substrate using KW and Wilcoxon rank-sum test (α= 0.05), respectively.

#### 2.5.3. Sequence Analysis

We used the SILVA project’s amplicon analysis pipeline (SILVAngs 1.4) [25] to analyze a dataset of 14 samples of sequence data. Within each sample reads were aligned to the SILVA SSU rRNA SEED using the SILVA Incremental Aligner (SINA v1.2.10 for ARB SVN (revision 21008)) [26] and subjected to quality controlled [27]. Reads shorter than 50 aligned nucleotides and those with more than 2 % of ambiguities or homopolymers were excluded, and putative contamination and artifacts, were identified and removed based on a low alignment quality (50 alignment identity, 40 alignment score reported by SINA); 702 of 1,401,269 reads were rejected. After quality control, identical reads were identified (dereplication), the unique reads were clustered (OTUs) on a per-sample basis. The reference read of each OTU was classified using using BLASTn (2.2.30+) [28] with the non-redundant version of the SILVA SSU Ref dataset v132 as the classification reference. Dereplication and clustering processes were performed using VSEARCH (version 2.14.2) [29], applying identity criteria of 1.00 ND 0.98, respectively. The classification of each OTU reference read was mapped onto all reads that were assigned to the respective OTU, providing quantitative information within the limitations of PCR and sequencing technique biases and multiple rRNA operons. Reads without any or weak classifications, where the function (% sequence identity + % alignment coverage)/2” did not exceed the value of 93, remained unclassified and were assigned to the meta-group “No Relative” as recommended by previous studies [30]. The data were analyzed with R version 3.6.1 [31] using RStudio (Version 1.2.1335 © 2009-2019 RStudio, Inc.). We used the relative abundance to observe qualitative differences in the taxonomic composition of the EcoPlates™ microbial communities between each treatment (BAC dosage in exposed and non-exposed AS). To visualize data, the ggplot2 package (cran.r-project.org) was used.

## 3. Results

### 3.1. BAC quantification and physicochemical parameters

The BAC concentration in the enriched AS reached 14.20 mg/L, which decreased to 2.27 mg/L after 96 hours.The control sample (not enriched) contained 3.14 mg/L of BAC after 96h. The pH was 7.1; dissolved oxygen 0.9 mg/L with a saturation of 2.1%, conductivity 580 μS/cm, and the temperature was 26.2 °C at the AS tank [32].

### 3.2. Multifunctionality index (MF) & Functional Diversity (FD)

The number of substrates used in treatment and control AS decreased as the BAC concentration on the EcoPlate™ increased (Table S1). This trend was more pronounced when the AUC threshold was increased, defining the substrate usage. The average number of substrates used was lower in the Control AS for all thresholds (Figure S1) and considering all thresholds together (Figure S2). However, differences between the Control and BAC (considering all BAC concentrations on EcoPlate™) were only statistically significant using T ≥ 0.5 (KW, p>0.05, Figure S3). The KW test followed by multiple comparisons with the Dunn test revealed significant differences between EcoPlate™ supplemented with 12.5 mg/L BAC and those with 50 mg/L BAC when T=0.1 or T=0.5 were used in control-treated and BAC-treated AS samples. These differences indicated a significant reduction in the number of substrates utilized by the exposed microbial community. However, no significant differences were observed in BAC supplementation on the EcoPlateTM when T=0.9 was used, either in treated or non-treated AS. Nevertheless, the BAC concentration supplemented on the EcoPlate™ had a stronger impact on FD (PERMANOVA, p=0.001). In this regard, differences in FD profile were observed after supplementing 50mg/L or more on the EcoPlate™ in both control and BAC-treated samples (Figure 2, PERMANOVA, p=0.002 and 0.004, respectively).

**Figure 2.**
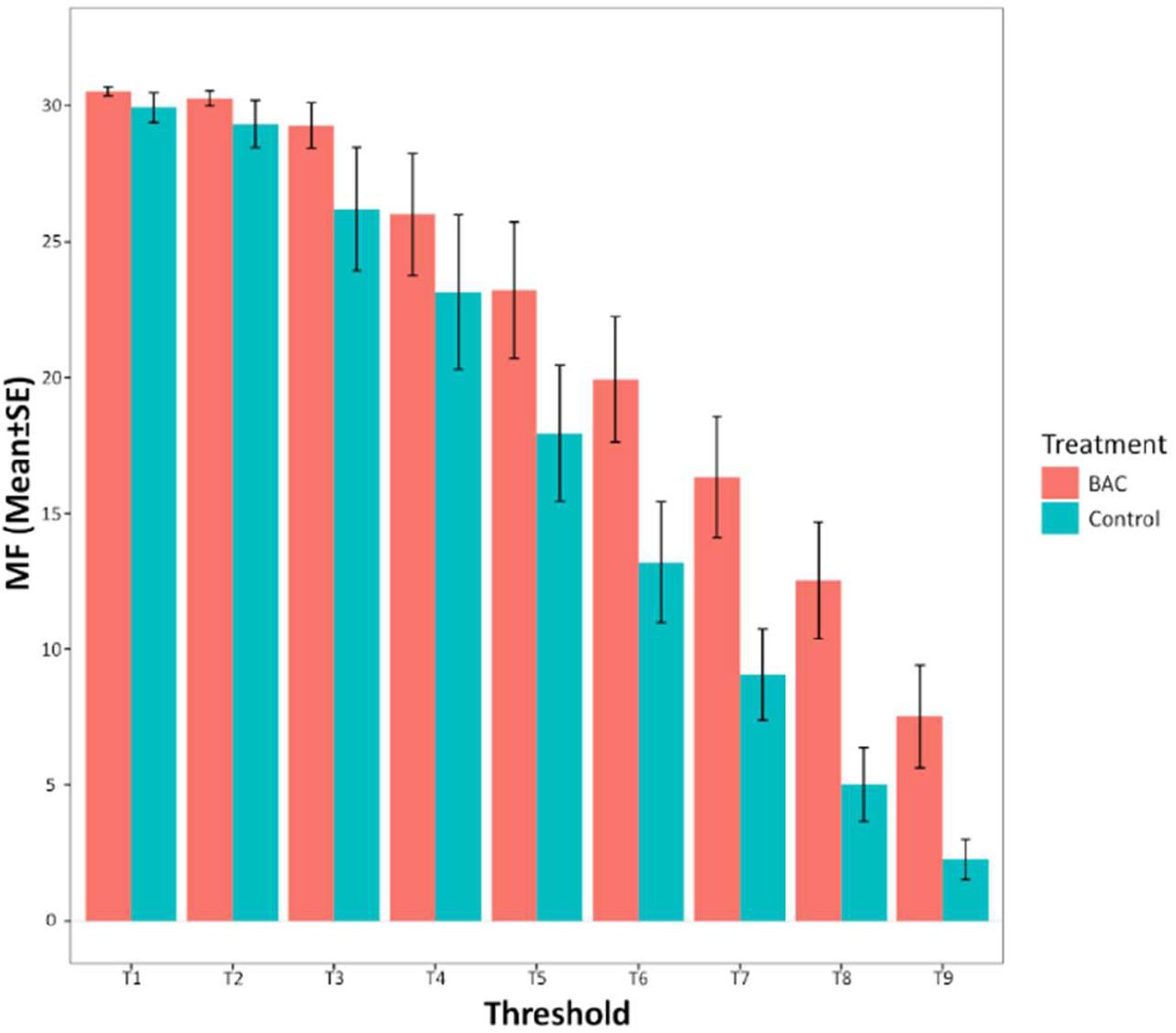
NMDS plots depict microbial functional diversity of Control (C), and BAC treated AS (B) based on dissimilarity matrixes by Bray Curtis Method using the AUC derived from the AUC measurements of each EcoPlate™.

### 3.3. Tolerance of microbial communities to BAC

Based on the average AUC from each concentration per replicate, BAC-treated ASs demonstrated a higher tolerance, with an EC_50_ of 57.8 mg/L (±sd=2.9) of BAC; while, the control AS showed a tolerance of 33.8 mg/L (±sd=12.8) of BAC. However, this difference was not statistically significant between the two treatments (KW, p= 0.12). For five substrates, the respiration measured in the control AS did not result in a dose-response curve from which an EC_50_ could be derivated, while this occurred with eight substrates for BAC-treated AS. In most of these cases, low or no inhibition was observed with increasing concentrations of BAC, and in few cases, there was even an increase in activity. When analyzing each substrate, the tolerance to increasing concentrations of BAC during respiration of polymer Tween40 was higher in control than on BAC-treated AS (Wilcoxon test, p<0.05). On the other hand, tolerance to BAC during respiration of carbohydrate β-Methyl-d-Glucoside and amino acid Glycyl-L-Glutamic Acid was higher in BAC-treated ASs (Wilcoxon test, p<0.05). For the rest of the substrates, no difference could be estimated at p=0.05 (Figure S4).

### 3.4. Phylogenetic analysis

After the assay, the community in the EcoPlates™ was mainly composed of the Gammaproteobacteria and Bacteroidetes phyla. It is important to mention that the composition that the AS supplemented with BAC presented more Bacteroidetes than control AS. The microbial community of the AS exposed to BAC underwent a composition shift at a concentration of 50 mg/L BAC, while the shift ocurred in the control microbial community at a lower BAC concentration (25 mg/L). Exposure to BAC on the EcoPlates™ caused Gammaproteobacteria to increase by over 80% and Bacteroidetes to decrease to less than 10 % of the global microbial composition in both communities. *Pseudomonas* was the main genus present in all samples from the Biolog® plates (Figure 3). The results showed an increase in the relative abundance of *Pseudomonas* and *Achromobacter* as the concentration of BAC increased; however, the changes in their abundance were more significant in dosed AS than in control AS. Other groups, such as *Elizabethkingia, Stenotrophomonas, Soonwooa*, and *Enterobacter*, were abundant in plates with low BAC concentration but diminished with concentrations around 12.5 mg/L of BAC (Figure 4).

**Figure 3.**
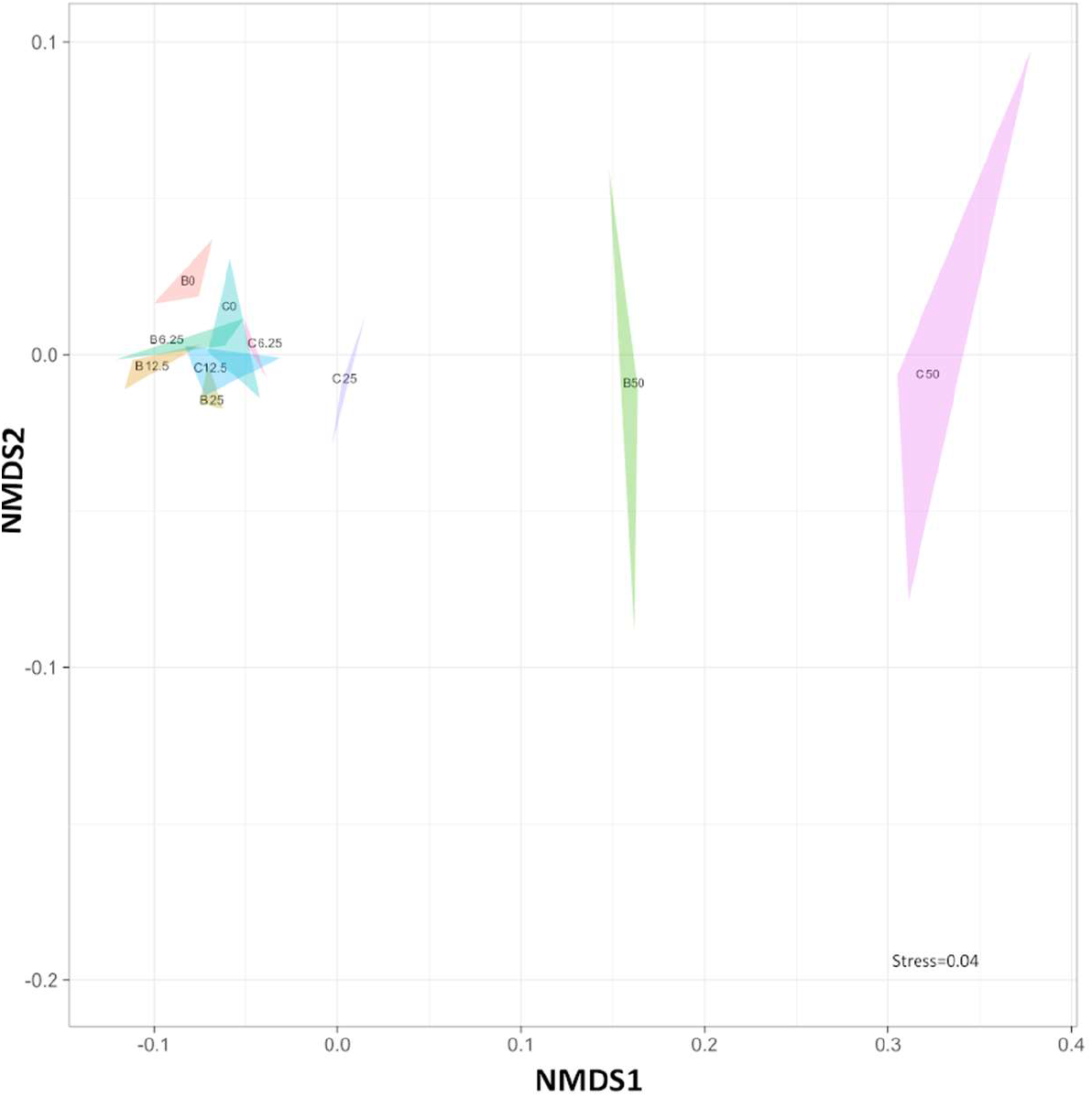
Phylogenetic composition of EcoPlates™ microbial suspensions after physiological profile assay. C=controls, D=dosed.

**Figure 4.**
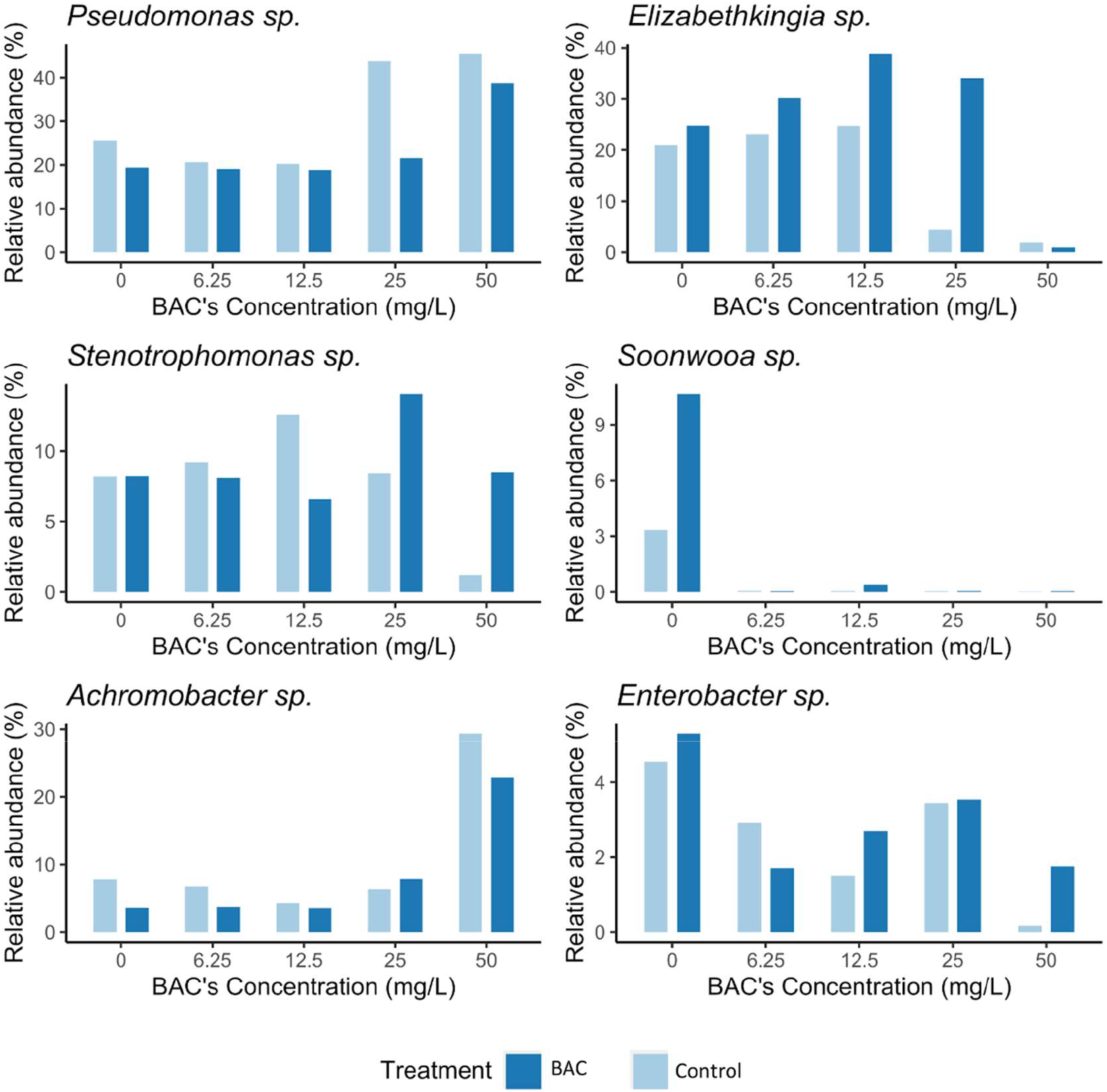
Main microbial genera present in EcoPlates™ microbial suspensions after physiological profile assay. A. *Pseudomonas*, B. *Elizabethkingia*, C. *Stenotrophomonas*, D. *Soonwooa*, E. *Achromobacter*, F. *Enterobacter*

## 4. Discussion

In this study, we demonstrated the rapid metabolic adaptation of the microbial community of AS from a household WWTP to a pulse of BAC at concentrations of mg per L. Our results suggest that such concentrations are a likely scenario in the WWTP in Costa Rica, as the functional diversity of the control microbial communities showed a similar magnitude of adaptation to BAC. The magnitude of the observed tolerance highlights the need to study the mechanisms of bacterial adaptation at the community level to antimicrobials, which are being used and detected in high quantities, such as quaternary ammonium compounds, in environmental samples.

BAC is constantly discharged into effluents from houses, hospitals, and industries as a component of formulated disinfectant products. Its use has also increased during the COVID-19 pandemic to eliminate the SARS-CoV-2 virus [33]. Recent reviews suggest that the concentrations of BAC entering wastewater treatment facility could oscillate between 0.29 to 28.8 μg/L [34]. However, exposure can reach up to ten times these concentrations [35], as analyzed in the residential WWTP in this study [36]. Therefore, the exposure scenario of BAC indicates a strong potential to exert a significant influence over the evolution of environmental bacteria, both genetically and phenotypically, as our results indicate.

In this experiment, the microbial community of the AS maintained its metabolic profile, as determined by Biolog® EcoPlates™, despite the high presence of antimicrobials in their environment. This was true for overall community metabolic diversity (FD), the number of substrates used (MF), and tolerance estimations (EC_50_) both in general, and for most of the carbon substrates individually. The adaptation in concentrations below 50mg/L of BAC was strongly evident. However, iIn EcoPlates™ with 50mg/L of BAC, the selection of highly tolerant groups during the *in vitro* 10mg/L nominal exposure of the AS seemed to have created deviations in the functional responses.

The physiological adaptation of AS microbial communities to different QAC is an increasing important area of research, as these compounds are known to accumulate in WAS treatment systems, potentially impacting the performance of WWTPs or WAS processing prior to final disposal. Chen et al. [37] previously demonstrated how increased BAC concentrations can affect the organic matter degradation, potentially altering the performance of wastewater treatment technologies by changing the organic matter content of the system. Microbial community tolerance to QACs has also been identified as an important parameter in modeling the ability of various WWTPs to function under different QACs concentrations [38].

The taxonomic analysis conducted to identify the microbial diversity of samplesin the EcoPlates™ reveled a significant proportion of adapted bacteria belonging to Gammaproteobacteria and Bacteroidetes phyla. The selection of r-strategists, fast-growing bacteria that dominate in nutrient-rich habitats, in EcoPlates™, belongs to Gammaproteobacteria (e.g., *Pseudomonas*), as observed in our study, has been reported before [39]. Notably, a high abundance of Proteobacteria and Bacteroidetes has also been described in WAS from municipal WWTPs [40]. Therefore, the main phyla detected on the EcoPlates™ correspond to the main groups identified previously. These findings confirm the usefulness of the metabolic profiling techniques in assessing AS. Although differences in the community composition at the genus level could be expected, the use of EcoPlates™ for physiological bioassays may contribute to these variations. Therefore, to ensure the accuracy of the results obtained using this metabolic profiling tool, further research is needed to address these differences in the community composition at genus level. We emphasize the importance of combining this type of phenotypical metabolic tool with the phylogenetic analysis of the communities represented in it.

In recent years, numerous studies have highlighted the potential of sublethal concentrations of BAC in increasing the tolerance and the antibiotic resistance profile of certain microbial community members in various environments. For instance, Kim et al. [41] reported that using BAC in a reactor with river sediment resulted in increased tolerance of the overall microbial community to 1,600 mg/L of BAC and modified the antibiotic resistance pattern in some bacterial isolates, remarkably increasing the resistance in *P. aeruginosa*.

The selection of Gammaproteobacteria, particularly of *Pseudomonas*, in the EcoPlates™ during our study supports the role of these genera in the community’s tolerance to BAC. The resistance to antimicrobial solutions containing BAC has been experimentally assessed in clinical isolates of *Pseudomonas* spp. since the 20^th^ century [42]. In a previous study we observed an increase in genes related to efflux pumps (*qacE/qacEΔ1*) after a BAC’s short-term exposure of AS to BAC [43]. This finding partially explains the observed tolerance to BAC, as other mechanisms may be ivolved, such as structural alterations in membranes [44].

The response to BAC exposure of the other bacterial genera identified in our study, namely *Achromobacter sp*., *Stenotrophomonas sp*., and *Elizabethkingia sp*., is consistent with previous reports in the literature. *Achromobacter xylosoxidans* has been shown to develop resistance after just one minute of exposure using a biofilm model with contact lenses [45]. *Stenotrophomonas maltophila* has been found to persist in hospital environments where BAC-containing disinfectants are intensively used, and its adaptation to BAC exposure has been associated with the activation of efflux pumps [46]. For *Elizabethkingia*, Forbes et al. [47] reported limited BAC tolerance with exposure to formulations containing it up to 0.1% (wt/vol) over a period of months in isolates of opportunistic pathogens from this genus in household environments.

## 5. Conclusion

The findings of this study indicate that the microbial community of the residential WWTP exhibits a high tolerance to BAC, as revealed by the metabolic profiling experiment using the Ecoplates system. This situation could be representative of other high-exposure settings, such as hospitals and agro-industrial wastewaters, where QACs are extensively used. The adaptation to QACs may be linked to the acquisition of antimicrobial-related resistance genes in certain bacterial groups. Given the significant exposure to QACs in terms of frequency and concentration, these compounds may act as a strong selective forces for aquatic microbial communities. However, further studies are required to better comprehend the impact of BAC selective force on the fitness and metabolism of bacteria at the community level in wastewater treatment plants.

Our results underscore the importance of examining the effects of BAC on wastewater and AS treatment, as changes in the community-level metabolism of carbon and nitrogen may also affect the quality of the effluents. Furthermore, future research should investigate the impact of introducing microbial groups adapted to high concentrations of BAC in the microbial activity of receiving water bodies.

## Nonstandard abbreviations

AS: activated sludge
AUC: area under the curve
BAC: benzalkonium chloride
EC_50_: half maximal effective concentration
FD: functional diversity
KW: Kruskal-Wallis test
MF: multifunctionality
OTU: operational taxonomic unit
PDA: Photodiode Array Detector
QAC: quaternary ammonium compound
Q-ToF: Quadrupole Time of Flight
UPLA: Ultra Performance Liquid Chromatography
WWTP: wastewater treatment plant

## 6. Acknowledgments

We thank Freylan Mena for its Lab support. And we also grateful to Consejo Nacional de Rectores (CONARE), Vicerrectoría de Investigación of Universidad Nacional, and Vicerrectoría de Investigación of Universidad de Costa Rica for funding this research. The authors have declared no conflict of interest.

