## Supplementary material for "Bacterial communities in residential wastewater treatment plants are physiologically adapted to high concentrations of quaternary ammonium compounds": Suplemental

### Supplementary Figures and Tables

Table S1. Number of substrates used (MF) for samples using quantile method thresholds. "C" and "B" indicate Control and BAC treatments respectively (replicates 1,2 and 3). The number after the hyphen is the BAC concentration in the EcoPlate.

* Indicates the thresholds for which statistical significance was found between Treatments.

| Sample | T=0.1 | T=0.2 | T=0.3 | T=0.4 | T=0.5 | T=0.6 | T=0.7 | T=0.8* | T=0.9* |
|---|---|---|---|---|---|---|---|---|---|
| C1_0 | 31 | 31 | 31 | 28 | 23 | 17 | 9 | 4 | 2 |
| C1_6.25 | 31 | 31 | 31 | 27 | 19 | 14 | 8 | 5 | 0 |
| C1_12.5 | 31 | 31 | 31 | 29 | 21 | 14 | 8 | 2 | 1 |
| C1_25 | 31 | 31 | 31 | 28 | 17 | 7 | 5 | 4 | 4 |
| C1_50 | 29 | 28 | 9 | 3 | 2 | 1 | 1 | 0 | 0 |
| C2_0 | 30 | 30 | 29 | 28 | 24 | 23 | 17 | 9 | 4 |
| C2_6.25 | 31 | 31 | 31 | 31 | 28 | 18 | 12 | 3 | 1 |
| C2_12.5 | 31 | 31 | 31 | 30 | 28 | 24 | 20 | 12 | 8 |
| C2_25 | 28 | 28 | 27 | 23 | 15 | 8 | 6 | 3 | 1 |
| C2_50 | 30 | 27 | 14 | 3 | 2 | 2 | 1 | 0 | 0 |
| C3_0 | 31 | 31 | 31 | 30 | 23 | 20 | 14 | 8 | 5 |
| C3_6.25 | 31 | 31 | 31 | 31 | 24 | 17 | 10 | 6 | 0 |
| C3_12.5 | 31 | 31 | 30 | 30 | 30 | 26 | 20 | 19 | 8 |
| C3_25 | 30 | 30 | 30 | 25 | 12 | 7 | 5 | 0 | 0 |
| C3_50 | 23 | 18 | 6 | 1 | 1 | 0 | 0 | 0 | 0 |
| B1_0 | 31 | 31 | 30 | 29 | 27 | 23 | 18 | 12 | 6 |
| B1_6.25 | 31 | 31 | 31 | 31 | 24 | 17 | 10 | 6 | 0 |
| B1_12.5 | 31 | 31 | 31 | 31 | 30 | 29 | 28 | 26 | 20 |

| | | | | | | | | | |
|---|---|---|---|---|---|---|---|---|---|
| B1_25 | 30 | 30 | 30 | 29 | 27 | 20 | 18 | 13 | 4 |
| B1_50 | 29 | 27 | 19 | 10 | 5 | 4 | 1 | 1 | 0 |
| B2_0 | 31 | 31 | 31 | 30 | 28 | 25 | 25 | 21 | 13 |
| B2_6.25 | 30 | 30 | 30 | 30 | 30 | 29 | 26 | 26 | 20 |
| B2_12.5 | 31 | 31 | 31 | 30 | 27 | 27 | 26 | 24 | 21 |
| B2_25 | 31 | 31 | 31 | 31 | 27 | 22 | 17 | 12 | 6 |
| B2_50 | 30 | 30 | 28 | 9 | 5 | 5 | 3 | 3 | 1 |
| B3_0 | 31 | 30 | 30 | 29 | 22 | 19 | 17 | 12 | 6 |
| B3_6.25 | 30 | 30 | 30 | 30 | 29 | 24 | 16 | 6 | 3 |
| B3_12.5 | 31 | 31 | 31 | 31 | 31 | 28 | 20 | 12 | 5 |
| B3_25 | 31 | 31 | 31 | 31 | 31 | 24 | 17 | 11 | 5 |
| B3_50 | 30 | 29 | 25 | 9 | 5 | 3 | 3 | 3 | 3 |

Figure S1. Average MF per threshold for all samples (0 to 50mg/L BAC on EcoPlate™) of Control and BAC Treatments. Bars indicate standard deviation.

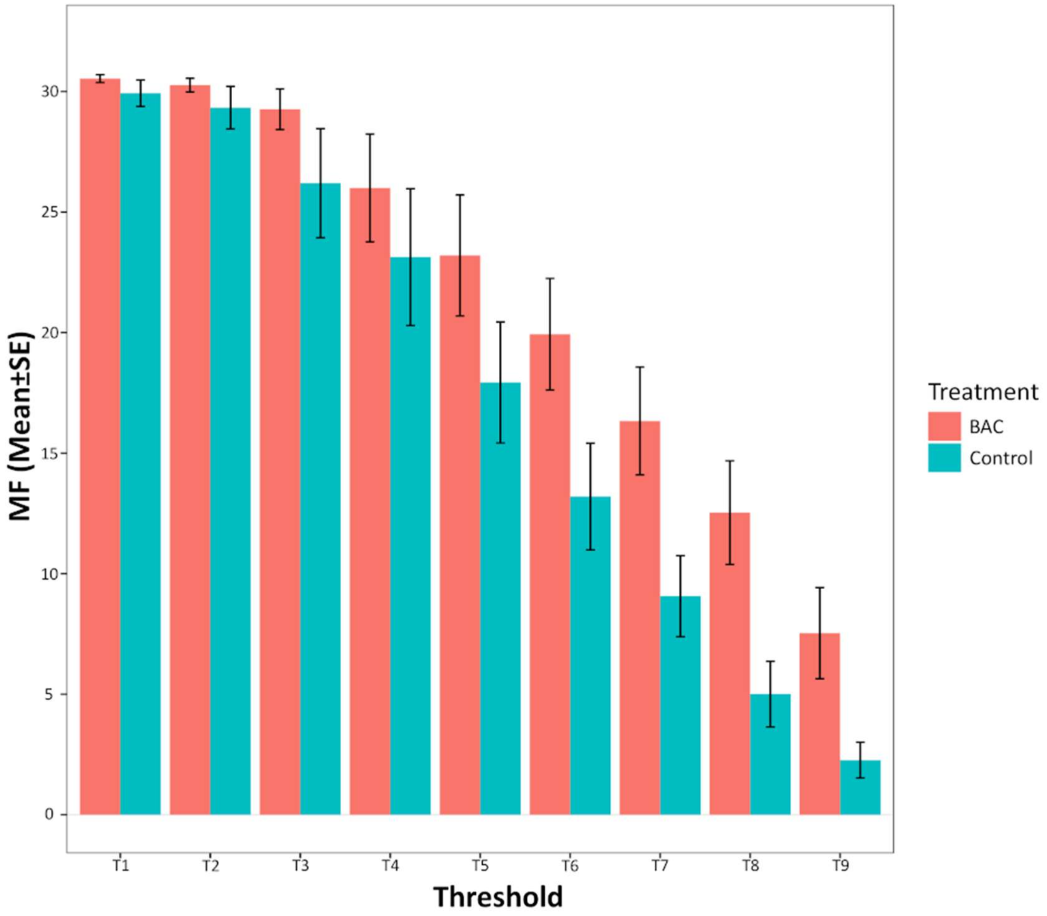

Figure S2. Average ±sd MF index for Control and BAC

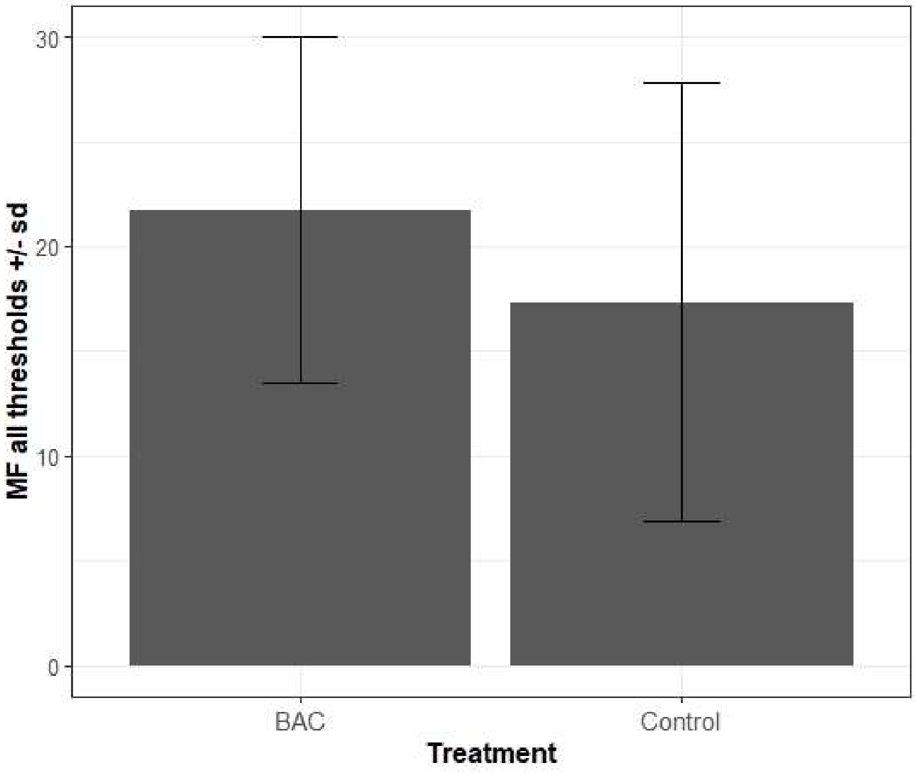

Figure S3. Average ±sd MF index for Control and BAC-pretreated waste activated sludge by threshold.

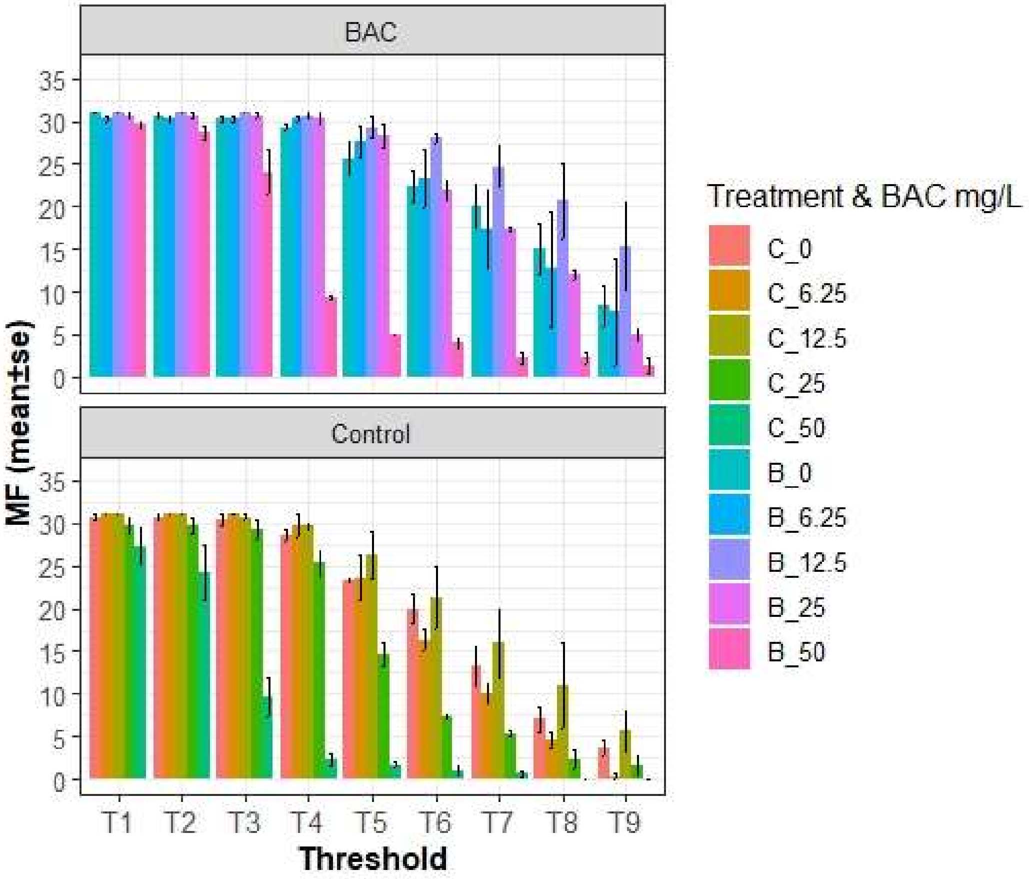

Figure S4. Mean $EC_{50}$ (n=3) per substrate in the EcoPlate$^{TM}$ (values represent mean BAC concentration in mg/L ± sd) for Control and BAC-treated ASs.

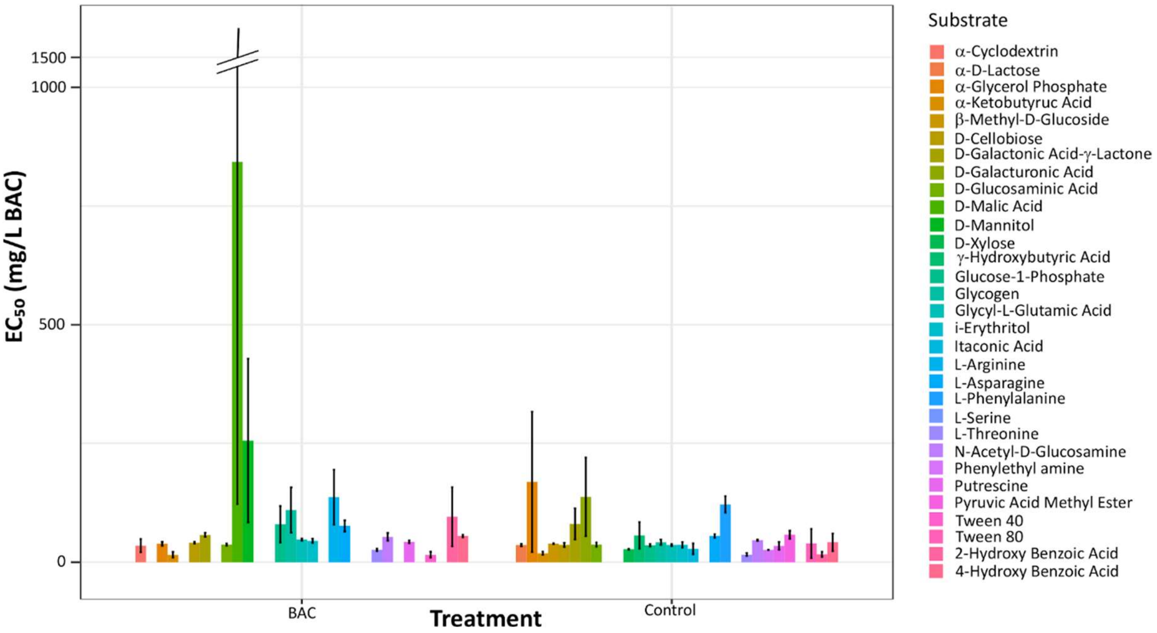
